# Transfer learning for biomedical named entity recognition with neural networks

**DOI:** 10.1101/262790

**Authors:** John M Giorgi, Gary D Bader

**Affiliations:** Department of Computer Science, University of Toronto, 10 Kings College Road, Toronto, Canada M5S 3G4; The Donnelly Centre, University of Toronto, 160 College Street, Toronto, Canada M5S 3E1; Department of Molecular Genetics, University of Toronto, 1 King’s College Circle, Toronto ON M5S 1A8

## Abstract

**Motivation:** The explosive increase of biomedical literature has made information extraction an increasingly important tool for biomedical research. A fundamental task is the recognition of biomedical named entities in text (BNER) such as genes/proteins, diseases, and species. Recently, a domain-independent method based on deep learning and statistical word embeddings, called long short-term memory network-conditional random field (LSTM-CRF), has been shown to outperform state-of-the-art entity-specific BNER tools. However, this method is dependent on gold-standard corpora (GSCs) consisting of hand-labeled entities, which tend to be small but highly reliable. An alternative to GSCs are silver-standard corpora (SSCs), which are generated by harmonizing the annotations made by several automatic annotation systems. SSCs typically contain more noise than GSCs but have the advantage of containing many more training examples. Ideally, these corpora could be combined to achieve the benefits of both, which is an opportunity for transfer learning. In this work, we analyze to what extent transfer learning improves upon state-of-the-art results for BNER.

**Results:** We demonstrate that transferring a deep neural network (DNN) trained on a large, noisy SSC to a smaller, but more reliable GSC significantly improves upon state-of-the-art results for BNER. Compared to a state-of-the-art baseline evaluated on 23 GSCs covering four different entity classes, transfer learning results in an average reduction in error of approximately 11%. We found transfer learning to be especially beneficial for target data sets with a small number of labels (approximately 6000 or less).

**Availability and implementation:** Source code for the LSTM-CRF is available at https://github.com/Franck-Dernoncourt/NeuroNER/ and links to the corpora are available at https://github.com/BaderLab/Transfer-Learning-BNER-Bioinformatics-2018/.

**Supplementary information:** Supplementary data are available at *Bioinformatics* online.

## 1 Introduction

The large quantity of biological information deposited in literature every day leads to information overload for biomedical researchers. In 2016 alone, there were 869,666 citations indexed in MEDLINE (https://www.nlm.nih.gov/bsd/index_stats_comp.html), which is greater than one paper per minute. Ideally, efficient, accurate text-mining and information extraction tools and methods could be used to help unlock structured information from this growing amount of raw text for use in computational data analysis. Text-mining has already proven useful for many types of large-scale biomedical data analysis, such as network biology (Zhou *et al.*, 2014; Al-Aamri *et al.*, 2017), gene prioritization (Aerts *et al.*, 2006), drug repositioning (Wang and Zhang, 2013; Rastegar-Mojarad *et al.*, 2015), and the creation of curated databases (Li *et al.*, 2015). A fundamental task in biomedical information extraction is the recognition of biomedical named entities in text (biomedical named entity recognition, BNER) such as genes and gene products, diseases, and species. Biomedical named entities have several characteristics that make their recognition in text particularly challenging (Campos *et al.*, 2012), including the sharing of head nouns (e.g. “91 and 84 kDa proteins” refers to “91 kDa protein” and “84 kDa protein”), several spelling forms per entity (e.g. “N-acetylcysteine”, “N-acetyl-cysteine”, and “NAcetylCysteine”) and ambiguous abbreviations (e.g. “TCF” may refer to “T cell factor” or to “Tissue Culture Fluid”). Until recently, state-of-the-art BNER tools have relied on hand-crafted features to capture the characteristics of different entity classes. This process of feature engineering, i.e. finding the set of features that best helps discern entities of a specific type from other tokens (or other entity classes), incurs extensive trial-and-error experiments. On top of this costly process, high-quality BNER tools typically employ entity-specific modules, such as whitelist and blacklist dictionaries, which are difficult to build and maintain. Defining these steps currently takes the majority of time and cost when developing BNER tools (Leser and Hakenberg, 2005) and leads to highly specialized solutions that cannot be ported to domains (or even entity types) other than the ones they were designed for. Very recently, however, a domain-independent method based on deep learning and statistical word embeddings, called long short-term memory network-conditional random field (LSTM-CRF), has been shown to outperform state-of-the-art entity-specific BNER tools (Habibi *et al.*, 2017). However, supervised, deep neural network (DNN) based approaches to BNER depend on large amounts of high quality, manually annotated data in the form of gold standard corpora (GSCs). The creation of a GSC is laborious: annotation guidelines must be established, domain experts must be trained, the annotation process is time-consuming and annotation disagreements must be resolved. As a consequence, GSCs in the biomedical domain tend to be small and focus on specific subdomains.

Silver-standard corpora (SSCs) represent an alternative that tend to be much larger, but of lower quality. SSCs are generated by using multiple, existing named entity taggers to annotate a large, unlabeled corpus. The heterogeneous results are automatically integrated, yielding a consensus-based, machine-generated ground truth. Compared to the generation of GSCs, this procedure is inexpensive, fast, and results in very large training datasets. The Collaborative Annotation of a Large Biomedical Corpus (CALBC) project sought to replace GSCs with SSCs, which would be much larger, more broadly scoped and more diversely annotated (Rebholz-Schuhmann *et al.*, 2010). However, Chowdhury and Lavelli (2011) found that a gene name recognition system trained on an initial version of the CALBC SSC performed worse than when trained on a BioCreative GSC. While SSCs have not proven to be viable replacements for GSCs, at least for the task of BNER, they do have the advantage of containing many more training examples (often in excess of 100 times more). This presents a unique transfer learning opportunity.

Transfer learning aims to perform a task on a “target” dataset using knowledge learned from a “source” dataset (Pan and Yang, 2010; Li, 2012; Weiss *et al.*, 2016). For DNNs, transfer learning is typically implemented by using some or all of the learned parameters of a DNN pre-trained on a source dataset to initialize training for a second DNN to be trained on a target dataset. Ideally, transfer learning improves generalization of the model, reduces training times on the target dataset, and reduces the amount of labeled data needed to obtain high performance. The idea has been successfully applied to many fields, such as speech recognition (Wang and Zheng, 2015), finance (Stamate *et al.*, 2015) and computer vision (Zeiler and Fergus, 2013; Yosinski *et al.*, 2014; Oquab *et al.*, 2014). Despite its popularity, few studies have been performed on transfer learning for DNN-based models in the field of natural language processing (NLP). For example, Mou *et al.* (2016) focused on transfer learning with convolutional neural networks (CNN) for sentence classification. To the best of our knowledge, there exists only one study which has analyzed transfer learning for DNN-based models in the context of NER (Lee *et al.*, 2017), and no study which has analyzed transfer learning for DNN-based approaches to BNER.

In this work, we analyze to what extent transfer learning on a source SSC to a target GSC improves performance on GSCs covering four different biomedical entity classes: chemicals, diseases, species and genes/proteins. We also identify the nature of these improvements and the scenarios where transfer learning offers the biggest advantages. The primary motivation for transfer learning from a SSC to a GSC is that we are able to expose the DNN to a large number of training examples (from the SSC) while minimizing the impact of noise in the SSC on model performance by also training on the GSC.

## 2 Materials and methods

The following sections present a technical explanation of the DNN architecture used in this study and in prior work (Habibi *et al.*, 2017). We first briefly describe LSTM, a specific kind of DNN, and then discuss the architecture of the hybrid LSTM-CRF model. We also describe the corpora used for evaluation and details regarding text pre-processing and evaluation metrics.

### 2.1 LSTM-CRF

Recurrent neural networks (RNNs) are popular for sequence labeling tasks, due to their ability to use previous information in a sequence for processing of current input. Although RNNs can, in theory, learn long-range dependencies, they fail to do so in practice and tend to be biased towards their most recent inputs in the sequence (Bengio *et al.*, 1994). An LSTM is a specific RNN architecture which mitigates this issue by keeping a memory cell that serves as a summary of the preceding elements of an input sequence and is able to model dependencies between sequence elements even if they are far apart (Hochreiter and Schmidhuber, 1997). The input to an LSTM unit is a sequence of vectors *x*_1_, *x*_2_, *…, x*_*T*_ of length *T*, for which it produces an output sequence of vectors *h*_1_, *h*_2_, *…h*_*T*_ of equal length by applying a non-linear transformation learned during the training phase. Each *h*_*t*_ is called the activation of the LSTM at token *t*, where a token is an instance of a sequence of characters in a document that are grouped together as a useful semantic unit for processing. The formula to compute one activation of an LSTM unit in the LSTM-CRF model is provided below (Lample *et al.*, 2016):

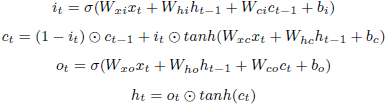

where all *W* s and *b*s are trainable parameters, *σ*(*·*) and *tanh*(*·*) denote the element-wise sigmoid and hyperbolic tangent activation functions, andʘ is the element-wise product. Such an LSTM-layer processes the input in one direction and thus can only encode dependencies on elements that came earlier in the sequence. As a remedy for this problem, another LSTM-layer which processes input in the reverse direction is commonly used, which allows detecting dependencies on elements later in the text. The resulting DNN is called a bi-directional LSTM (Graves and Schmidhuber, 2005). The representation of a word using this model is obtained by concatenating its left and right context representations, 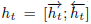 These representations effectively encode a representation of a word in context. Finally, a sequential conditional random field (Lafferty *et al.*, 2001) receives as input the scores output by the bi-directional LSTM to jointly model tagging decisions. LSTM-CRF (Lample *et al.*, 2016) is a domain-independent NER method which does not rely on any language-specific knowledge or resources, such as dictionaries. In this study, we used NeuroNER (Dernoncourt *et al.*, 2017b), a named entity recognizer based on a bi-directional LSTM-CRF architecture. The major components of the LSTM-CRF model for sequence tagging are described below:

1. **Token embedding layer** maps each token in the input sequence to a token embedding.
2. **Character embedding layer** maps each character in the input sequence to a character embedding.
3. **Character Bi-LSTM layer** takes as input character embeddings and outputs a single, character-level representation vector that summarizes the information from the sequence of characters in the corresponding token.
4. **Token Bi-LSTM layer** takes as input a sequence of character-enhanced token vectors, which are formed by concatenating the outputs of the token embedding layer and the character Bi-LSTM layer.
5. **Label prediction layer** using a fully-connected neural network, maps the output from the token Bi-LSTM layer to a sequence of vectors containing the probability of each label for each corresponding token.
6. **Label sequence optimization layer** using a CRF, outputs the most likely sequence of predicted labels based on the sequence of probability vectors from the previous layer.

Figure 1 illustrates the DNN architecture. All layers of the network are learned jointly. A detailed description of the architecture is explained in Dernoncourt *et al.* (2017a).

**Fig. 1.**
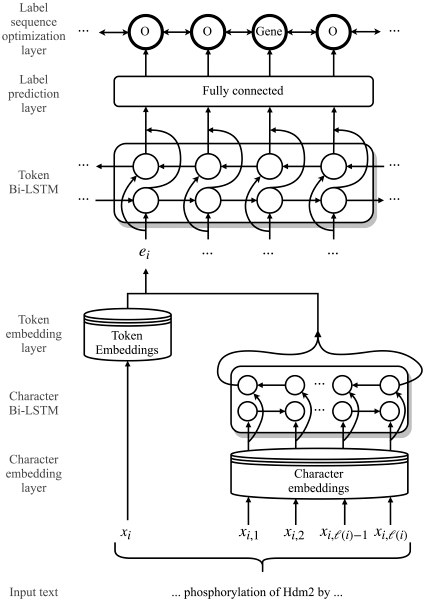
Architecture of the hybrid long short-term memory network-conditional random field (LSTM-CRF) model for named entity recognition (NER). Here, *x*_*i*_ is the *i*^*th*^ token in the input sequence, *x*_*ij*_ is the *j*^*th*^ character of the *i*^*th*^ token, *£*(*i*) is the number of characters in the *i*^*th*^ token and *ei* is the character-enhanced token embedding of the *i*^*th*^ token. For transfer learning experiments, we train the parameters of the model on a source dataset, and transfer all of the parameters to initialize the model for training on a target dataset.

#### 2.1.1 Training

The network was trained using the back-propagation algorithm to update the parameters on every training example, one at a time, using stochastic gradient descent. For regularization, dropout was applied before the token Bi-LSTM layer, and early stopping was used on the validation set with a patience of 10 epochs, i.e., the model stopped training if performance did not improve on the validation set for 10 consecutive epochs. While training on the source datasets, the learning rate was set to 0.0005, gradient clipping to 5.0 and the dropout rate to 0.8. These hyperparameters were chosen to discourage convergence of the network on the source dataset, such that further learning could occur on the target dataset. While training on the target datasets, the learning rate was increased to 0.005, and the dropout rate lowered to 0.5. These are the default hyperparameters of NeuroNER and give good performance on most NER tasks. Additionally, Lample *et al.* (2016) showed a dropout rate of 0.5 to be optimal for the task of NER.

### 2.2 Gold standard corpora

We performed our evaluations on four entity types: chemicals, diseases, species and genes/proteins. We used 23 datasets (i.e., GSCs), each containing hand-labeled annotations for one of these entity types, such as the “CDR” corpus for chemicals (Li *et al.*, 2016), “NCBI Disease” for disease names (Doğan *et al.*, 2014), “S800” for species (Pafilis *et al.*, 2013) and “DECA” for genes/proteins (Wang *et al.*, 2010). Table 1 lists all corpora and their characteristics, like the number of sentences, tokens and annotated entities per entity class (measured after text pre-processing as described in Section 2.6).

**Table 1.**
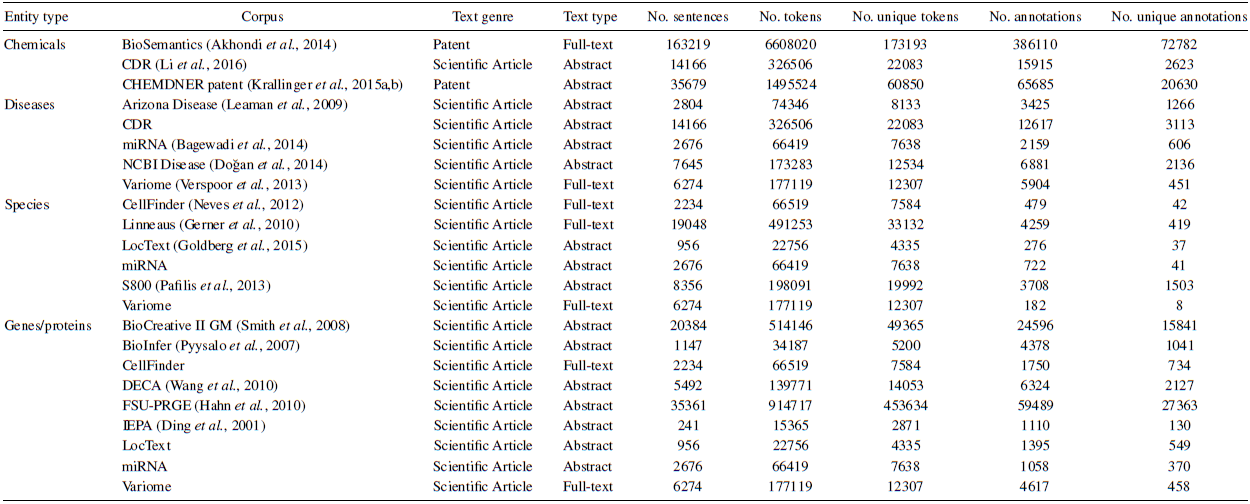
Gold standard corpora (GSCs) used in this work.

### 2.3 Silver standard corpora

We collected 50,000 abstracts (from a total of 174,999) at random from the CALBC-SSC-III-Small corpus (Kafkas *et al.*, 2012) for each of the four entity types it annotates: chemicals and drugs (*CHED*), diseases (*DISO*), living beings (*LIVB*), and, genes/proteins (*PRGE*). These SSCs served as the source datasets for each transfer learning experiment we performed. For each SSC, any document that appeared in at least one of the GSCs annotated for the same entity type was excluded to avoid possible circularity in performance testing (e.g., if a document with PubMed ID 130845 was found in a GSC annotated for genes/proteins, it was excluded from the PRGE SSC). In an effort to reduce noise in the SSCs, a selection of entities present but not annotated in any of the GSCs of the same entity type were removed from the SSCs. For example, certain text spans such as “genes”, “proteins”, and “animals” are annotated in the SSCs but not annotated in any of the GSCs of the same entity type, and so were removed from the SSCs (see Supplementary Material).

### 2.4 Word embeddings

We utilized statistical word embedding techniques to capture functional (i.e., semantic and syntactic) similarity of words based on their surrounding words. Word embeddings are pre-trained using large unlabeled datasets typically based on token co-occurrences (Collobert *et al.*, 2011; Mikolov *et al.*, 2013; Pennington *et al.*, 2014). The learned vectors, or word embeddings, encode many linguistic regularities and patterns, some of which can be represented as linear translations. In the canonical example, the resulting vector for vec(“king”) *-* vec(“man”) + vec(“woman”) is closest to the vector associated with “queen”, i.e., vec(“queen”). We used the *Wiki-PubMed-PMC* model, trained on a combination of PubMed abstracts (nearly 23 million abstracts) and PubMedCentral (PMC) articles (nearly 700,000 full-text articles) plus approximately four million English Wikipedia articles, and therefore mixes domain-specific texts with domain-independent ones. The model was created by Pyysalo *et al.* (2013) using Google’s word2vec (Mikolov *et al.*, 2013). We chose this model to be consistent with Habibi *et al.*, 2017, who showed it to be optimal for the task of BNER and who we compare to.

### 2.5 Character embeddings

The token-based word embeddings introduced above effectively capture distributional similarities of words (where does the word tend to occur in a corpus?) but are less effective at capturing orthographic similarities (what does the word look like?). In addition, token-based word embeddings cannot account for out-of-vocabulary tokens and misspellings. Character-based word representation models (Ling *et al.*, 2015) offer a solution to these problems by using each individual character of a token to generate the token vector representation. Character-based word embeddings encode sub-token patterns such as morphemes (e.g. suffixes and prefixes), morphological inflections (e.g. number and tense) and other information not contained in the token-based word embeddings. The LSTM-CRF architecture used in this study combines character-based word representations with token-based word representations, allowing the model to learn distributional and orthographic features of words. Character embeddings are initialized randomly and learned jointly with the other parameters of the DNN.

### 2.6 Text pre-processing

All corpora were first converted to the Brat standoff format (http://brat.nlplab.org/standoff.html). In this format, annotations are stored separately from the annotated document text. Thus, for each text document in the corpus, there is a corresponding annotation file. The two files are associated by the file naming convention that their base name (file name without suffix) is the same. All annotations follow the same basic structure: each line contains one annotation, and each annotation has an identifier that appears first on the line, separated from the rest of the annotation by a single tab character.

### 2.7 Evaluation metrics

We randomly divided each GSC into three disjoint subsets. 60% of the samples were used for training, 10% as the development set for the training of methods, and 30% for the final evaluation. We compared all methods in terms of precision, recall, and F1-score on the test sets. Precision is computed as the percentage of predicted labels that are gold labels (i.e., labels that appear in the GSC), recall as the percentage of gold labels that are correctly predicted, and F1-score as the harmonic mean of precision and recall. A predicted label is considered correct if and only if it exactly matches a gold label. NeuroNER uses the conlleval script from the CoNLL-2000 shared task to compute all performance metrics (https://www.clips.uantwerpen.be/conll2000/chunking/output.html).

For a single performance metric across all corpora, we compute the average percent reduction in error, which in our case is the average reduction in F1-score error due to transfer learning (TL) relative to the baseline:

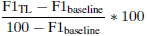

## 3 Results

We assessed the effect of transfer learning on the performance of a state-of-the-art method for BNER (LSTM-CRF) on 23 different datasets covering four different types of biomedical entity classes. We applied transfer learning by training all parameters of the DNN on a source dataset (CALBC-SSC-III) for a particular entity type (e.g. genes/proteins) and used the same DNN to retrain on a target dataset (i.e. a GSC) of the same entity type. Results were compared to training the model only on the target dataset using the same word embeddings (baseline).

### 3.1 Quantifying the impact of transfer learning

In this experiment, we determine whether transfer learning improves on state-of-the-art results for BNER. Table 2 compares the macro-averaged performance metrics of the model trained only on the target dataset (i.e., the baseline) against the model trained on the source dataset followed by the target dataset for 23 evaluation sets; exact precision, recall and F1-scores are given in Appendix A. Transfer learning improves the average F1-scores over the baseline for each of the four entity classes, leading to an average reduction in error of 11.28% across the GSCs. On corpora annotated for diseases, species, and genes/proteins, transfer learning (on average) improved both precision and recall, leading to sizable improvements in F1-score. For corpora annotated for chemicals, transfer learning (on average) slightly increased recall at the cost of precision for a small increase in F1-score. More generally, transfer learning appears to be especially effective on corpora with a small number of labels. For example, transfer learning led to a 9.69% improvement in F1-score on the test set of the CellFinder corpus annotated for genes/proteins — the eighth smallest corpus overall by number of labels. Conversely, the only GSC for which transfer learning worsened the performance compared to the baseline was the BioSemantics corpora, the largest GSC used in this study.

**Table 2.** Macro-averaged performance values in terms of precision, recall and F1-score for baseline (B) and transfer learning (TL) over the corpora per each entity type. Baseline values are derived from training on the target dataset only, while transfer learning values are derived by training on the source dataset followed by training on the target dataset. The macro average is computed by averaging the performance scores obtained by the classifiers for each corpus of a given entity class. Bold: best scores.

|  | Precision (%) |  | Recall (%) |  | F1-score (%) |  |
| --- | --- | --- | --- | --- | --- | --- |
|  | B | TL | B | TL | B | TL |
| Chemicals | **87.10** | 87.05 | 89.19 | **89.47** | 88.08 | **88.21** |
| Diseases | 80.41 | **81.41** | 81.13 | **82.46** | 80.73 | **82.09** |
| Species | 84.18 | **84.52** | 84.44 | **90.12** | 84.20 | **87.01** |
| Genes/proteins | 82.09 | **83.38** | 80.85 | **83.08** | 81.20 | **83.09** |

### 3.2 Learning curve for select evaluation sets

Figure 2 compares learning curves for the baseline model against the model trained with transfer learning on select GSCs, one for each entity class. The number of training examples used as the target training set is reported as a percent of the overall GSC size (e.g., for a GSC of 100 documents, a target train set size of 60% corresponds to 60 documents). The performance improvement due to transfer learning is especially pronounced when a small number of labels are used as the target training set. For example, on the miRNA corpus annotated for diseases, performing transfer learning and using 10% of examples as the train set leads to similar performance as using approximately 28% of examples as the train set when not using transfer learning. The performance gains from transfer learning diminish as the number of training examples used for the target training set increases.

**Fig. 2.**
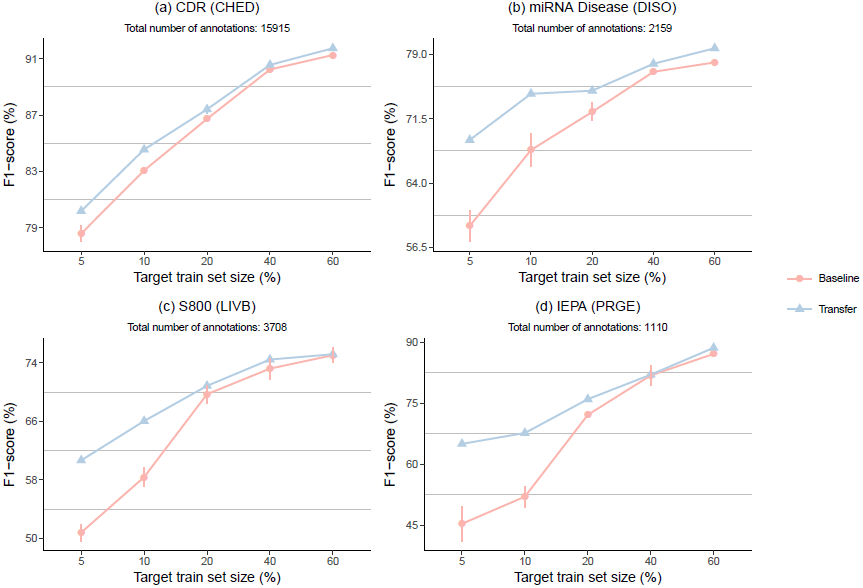
Impact of transfer learning on the F1-scores. Baseline corresponds to training the model only with the target dataset, and transfer learning corresponds to training on the source dataset followed by training on the target dataset. The number of training examples used in the target training set is reported as a percent of the overall GSC size (e.g., for a GSC of 100 documents, a target train set size of 60% corresponds to 60 documents). the size of the data point symbol.

These results suggest that transfer learning is especially beneficial for datasets with a small number of labels. Figure 3 more precisely captures this trend. Large improvements in F1-score are observed for corpora with up to approximately 6000 total annotations, with improvement quickly tailing off afterward. Indeed, all corpora for which transfer learning led to a statistically significant (*p ≤* 0.05) improvement in F1-score have 6000 annotations or less (see Appendix A). Therefore, it appears that the expected performance improvements derived from transfer learning are largest when the number of annotations in the target dataset is approximately 6000 or less.

**Fig. 3.**
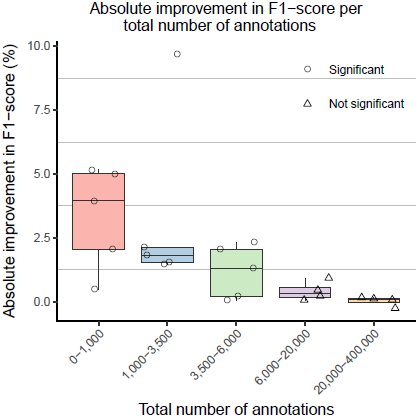
Box plots representing absolute F1-score improvement over the b transfer learning, grouped by the total number of annotations in the target g corpora (GSCs). Bin boundaries were generated using the R package binr (I Scores for individual GSCs are plotted, where point shapes indicate statistical (*p ≤* 0.05).

### 3.3 Error analysis

We compared the errors made by the baseline and transfer learning classifiers by computing intersections of true-positives (TPs), false-negatives (FNs) and false-positives (FPs) per entity type (Figure 4). In general, there is broad agreement for baseline and transfer learning classifiers, especially for TPs, with different strengths per entity type. For diseases, transfer learning has a negligible impact on the number of TPs and FNs but leads to a sizable decrease in the number of FPs, thereby increasing precision. For species and genes/proteins, transfer learning decreases the number of FNs but increases the number of FPs and TPs — trading precision for recall. Interestingly, the pattern of TPs, FPs, and FNs for chemical entities appears to disagree with the pattern in performance metrics observed at the macro-level (Table 2). However, because the decrease in FPs and TPs is roughly equal in magnitude to the increase in FNs, the net effect is the nearly identical performance metrics of the baseline and transfer learning method for chemical entities that we observe at the macro-level.

**Fig. 4.**
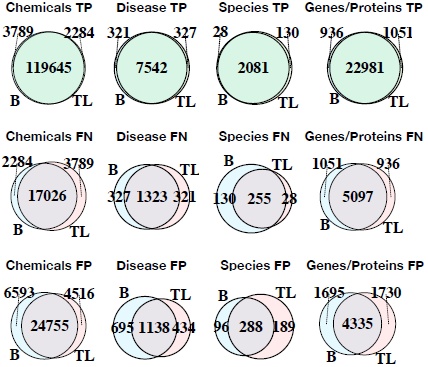
Venn diagrams demonstrating the area of overlap among the true-positive (TP), false-negative (FN), and false-positive (FP) sets of the baseline (B) and transfer learning (TL) methods per entity class.

## 4 Discussion

In this study, we demonstrated that transfer learning from large SSCs (source) to smaller GSCs (target) improves performance over training solely on the GSCs for BNER. This is the first study of the effect of transfer learning for BNER. On average, transfer learning leads to improvements in F1-score over a state-of-the-art baseline, especially for smaller GSCs, though the nature and degree of these improvements vary per entity type (Table 2).

The effect of transfer learning is most pronounced when the target train set size is small, with improvements diminishing as the training set size grows (Figure 2). Significant improvements in performance were observed only for corpora with 6000 total annotations or less (Figure 3). We conclude that the representations learned from the source dataset are effectively transferred and exploited for the target dataset; when transfer learning is adopted, fewer annotations are needed to achieve the same level of performance as when the source dataset is not used. Thus, our results suggest that researchers and text-mining practitioners can make use of transfer learning to reduce the number of hand-labeled annotations necessary to obtain high performance for BNER. We also suggest that transfer learning is likely to be a valuable tool for existing GSCs with a small number of labels.

Transfer learning had little impact on performance for chemical GSCs. This is likely explained by the much larger size of these corpora, which have a median number of annotations ten times that of the next largest set of corpora (diseases). We suspect that relatively large corpora contain enough training examples for the model to generalize well, in which case we would not expect transfer learning to improve model performance. Indeed, the largest corpora in our study, the BioSemantics corpora (annotated for chemical entities), was the only corpora for which transfer learning worsened performance over the baseline. With 386,110 total annotations (almost double the sum total of annotations in the remaining 22 GSCs) the BioSemantics corpus is an extreme outlier. To create such a large GSC, documents were first pre-annotated and made available to four independent annotator groups each consisting of two to ten annotators (Akhondi *et al.*, 2014). This is a much larger annotation effort than usual and is not realistic for the creation of typical GSCs in most contexts. Another possibility is that chemical entities are easier to identify in text.

BNER has recently made substantial advances in performance with the application of deep learning (Habibi *et al.*, 2017). We show that transfer learning is a valuable addition to this method. However, there are opportunities to further optimize this approach, for instance, by determining the optimal size of the source dataset, developing robust methods of filtering noise from the source dataset, and extensive hyperparameter tuning (Young *et al.*, 2015; Reimers and Gurevych, 2017).

### 4.1 Related work

Lee *et al.* (2017) performed a similar set of experiments, transferring an LSTM-CRF based NER model from a large labeled dataset to a smaller dataset for the task of de-identification of protected health information (PHI) from electronic health records (EHR). It was demonstrated that transfer learning improves the performance over state-of-the-art results, and may be especially beneficial for a target dataset with a small number of labels. Our results confirm these findings in the context of BNER. The study also explored the importance of each layer of the DNN in transfer learning. They found that transferring a few lower layers is almost as efficient as transferring all layers, which supports the common hypothesis that higher layers of DNN architectures contain the parameters that are more specific to the task and dataset used for training. We performed a similar experiment (see Supplementary Figure 1) with similar results.

A recent study has sought to adopt multi-task learning for the task of BNER (Crichton *et al.*, 2017). While transfer learning and multi-task learning are different, they are both forms of inductive transfer that are often employed for similar reasons. At a high-level, multi-task learning (Caruana, 1993) is a machine learning method in which multiple learning tasks are solved at the same time. This is in contrast to transfer learning, where we typically transfer some knowledge learned from one task or domain to another. In the classification context, multi-task learning is used to improve the performance of multiple classification tasks by learning them jointly. The idea is that by sharing representations between tasks, we can exploit commonalities, leading to improved learning efficiency and prediction accuracy for the task-specific models, when compared to training the models separately (Thrun, 1996; Caruana, 1998; Baxter *et al.*, 2000). Crichton *et al.* (2017) demonstrated that a neural network multi-task model outperforms a comparable single-task model, on average, for the task of BNER. Perhaps most interestingly, it was found that the performance improvements due to multi-task learning diminish as the size of the datasets grows – something we found to be true of transfer learning as well. Together, our results suggest that there is promise in the idea of sharing information between tasks and between datasets for biomedical text-mining, and may help overcome the limitations of training DNNs on small biomedical GSCs. Future work could evaluate the combination of multi-task and transfer learning to see if they are complementary and can further improve performance.

## 5 Conclusion

In this work, we have studied transfer learning with DNNs for BNER (specifically LSTM-CRF) by transferring parameters learned on large, noisy SSC for fine-tuning on smaller, but more reliable GSC. We demonstrated that compared to a state-of-the-art baseline evaluated on 23 GSCs, transfer learning results in an average reduction in error of approximately 11%. The largest performance improvements were observed for GSCs with a small number of labels (on the order of 6000 or less). Our results suggest that researchers and text-mining practitioners can make use of transfer learning to reduce the number of hand-labeled annotations necessary to obtain high performance for BNER. We also suggest that transfer learning is likely to be a valuable tool for existing GSC with a small number of labels. We hope this study will increase interest in the development of large, broadly-scoped SSCs for the training of supervised biomedical information extraction methods.

## Acknowledgements

We acknowledge the creators of the various corpora used in this study. We gratefully acknowledge the support of NVIDIA Corporation with the donation of the Titan Xp GPU used for this research.

## Appendix A

We have provided precision, recall, and F1-scores of each model (baseline and transfer learning) for each corpus in Table A1. We also include the scores reported by Habibi *et al.* (2017) for comparison. We measure statistical significance using a two-tailed t-test.

**Table 3.**
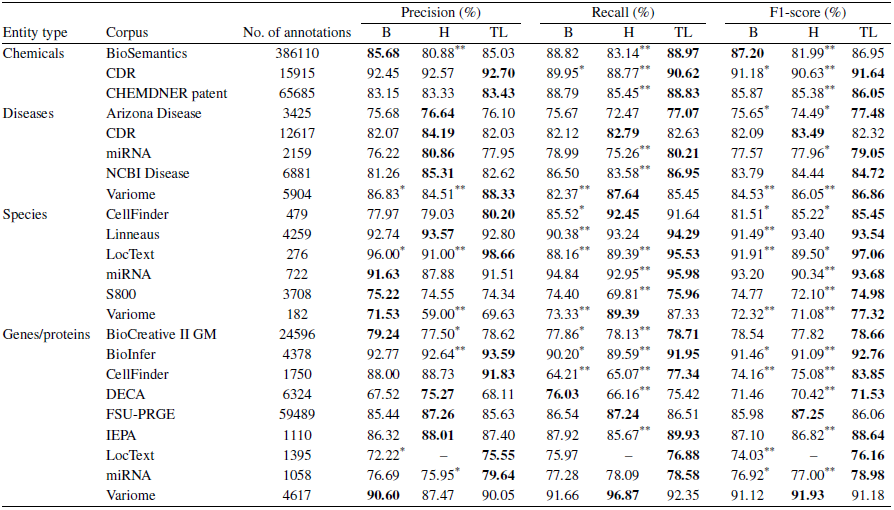
Performance values in terms of precision, recall and F1-score for our baseline (B) and transfer learning (TL) methods and those reported by Habibi et al. (2017) (H). Baseline values are derived from training on the target dataset only, while transfer learning values are derived by training on the source dataset followed by training on the target dataset. The baseline and transfer learning methods were trained for five runs, and the results were averaged. Scores from Habibi et al. (2017) are reported directly. Bold: best scores, *: significantly worse than the TL model (*p ≤* 0.05), **: significantly worse than the TL model (*p ≤* 0.01).

## References

Aerts, S., Lambrechts, D., Maity, S., Loo, P. V., Coessens, B., Smet, F. D., Tranchevent, L.-C., Moor, B. D., Marynen, P., Hassan, B., Carmeliet, P., and Moreau, Y. (2006). Gene prioritization through genomic data fusion. Nature Biotechnology, 24(5), 537–544.

Akhondi, S. A., Klenner, A. G., Tyrchan, C., Manchala, A. K., Boppana, K., Lowe, D., Zimmermann, M., Jagarlapudi, S. A., Sayle, R., Kors, J. A., et al. (2014). Annotated chemical patent corpus: a gold standard for text mining. PloS one, 9(9), e107477.

Al-Aamri, A., Taha, K., Al-Hammadi, Y., Maalouf, M., and Homouz, D. (2017). Constructing genetic networks using biomedical literature and rare event classification. Scientific reports, 7(1), 15784.

Bagewadi, S., Bobić, T., Hofmann-Apitius, M., Fluck, J., and Klinger, R. (2014). Detecting mirna mentions and relations in biomedical literature. F1000Research, 3.

Baxter, J. et al. (2000). A model of inductive bias learning. J. Artif. Intell. Res.(JAIR), 12(149-198), 3.

Bengio, Y., Simard, P., and Frasconi, P. (1994). Learning long-term dependencies with gradient descent is difficult. IEEE transactions on neural networks, 5(2), 157–166.

Campos, D., Matos, S., and Oliveira, J. L. (2012). Biomedical named entity recognition: A survey of machine-learning tools. In S. Sakurai, editor, Theory and Applications for Advanced Text Mining, chapter 08. InTech, Rijeka.

Caruana, R. (1993). Multitask learning: A knowledge-based source of inductive bias. In Proceedings of the Tenth International Conference on Machine Learning, pages 41–48. Citeseer.

Caruana, R. (1998). Multitask learning. In Learning to learn, pages 95–133. Springer.

Chowdhury, F. M. and Lavelli, A. (2011). Assessing the practical usability of an automatically annotated corpus. In Proceedings of the 5th Linguistic Annotation Workshop, pages 101–109. Association for Computational Linguistics.

Collobert, R., Weston, J., Bottou, L., Karlen, M., Kavukcuoglu, K., and Kuksa, P. (2011). Natural language processing (almost) from scratch. Journal of Machine Learning Research, 12(Aug), 2493–2537.

Crichton, G., Pyysalo, S., Chiu, B., and Korhonen, A. (2017). A neural network multi-task learning approach to biomedical named entity recognition. BMC Bioinformatics, 18(1), 368.

Dernoncourt, F., Lee, J. Y., Uzuner, O., and Szolovits, P. (2017a). De-identification of patient notes with recurrent neural networks. Journal of the American Medical Informatics Association, 24(3), 596–606.

Dernoncourt, F., Lee, J. Y., and Szolovits, P. (2017b). NeuroNER: an easy-to-use program for named-entity recognition based on neural networks. Conference on Empirical Methods on Natural Language Processing (EMNLP).

Ding, J., Berleant, D., Nettleton, D., and Wurtele, E. (2001). Mining medline: abstracts, sentences, or phrases? In Biocomputing 2002, pages 326–337. World Scientific.

Dogăn, R. I., Leaman, R., and Lu, Z. (2014). NCBI disease corpus: A resource for disease name recognition and concept normalization. Journal of Biomedical Informatics, 47, 1–10.

Gerner, M., Nenadic, G., and Bergman, C. M. (2010). Linnaeus: a species name identification system for biomedical literature. BMC bioinformatics, 11, 85.

Goldberg, T., Vinchurkar, S., Cejuela, J. M., Jensen, L. J., and Rost, B. (2015). Linked annotations: a middle ground for manual curation of biomedical databases and text corpora. BMC Proceedings, 9(5), A4.

Graves, A. and Schmidhuber, J. (2005). Framewise phoneme classification with bidirectional lstm and other neural network architectures. Neural Networks, 18(5), 602–610.

Habibi, M., Weber, L., Neves, M., Wiegandt, D. L., and Leser, U. (2017). Deep learning with word embeddings improves biomedical named entity recognition. Bioinformatics, 33(14), i37–i48.

Hahn, U., Tomanek, K., Beisswanger, E., and Faessler, E. (2010). A proposal for a configurable silver standard. In Proceedings of the Fourth Linguistic Annotation Workshop, LAW IV ’10, pages 235–242, Stroudsburg, PA, USA. Association for Computational Linguistics.

Hochreiter, S. and Schmidhuber, J. (1997). Long short-term memory. Neural computation, 9(8), 1735–1780.

Izrailev, S. (2015). binr: Cut Numeric Values into Evenly Distributed Groups. R package version 1.1.

Kafkas, S., Lewin, I., Milward, D., van Mulligen, E. M., Kors, J. A., Hahn, U., and Rebholz-Schuhmann, D. (2012). Calbc: Releasing the final corpora. In LREC, pages 2923–2926.

Krallinger, M., Rabal, O., Leitner, F., Vazquez, M., Salgado, D., Lu, Z., Leaman, R., Lu, Y., Ji, D., Lowe, D. M., et al. (2015a). The chemdner corpus of chemicals and drugs and its annotation principles. Journal of cheminformatics, 7(S1), S2.

Krallinger, M., Rabal, O., Lourenço, A., Perez, M. P., Rodriguez, G. P., Vazquez, M., Leitner, F., Oyarzabal, J., and Valencia, A. (2015b). Overview of the chemdner patents task. In Proceedings of the fifth BioCreative challenge evaluation workshop, pages 63–75.

Lafferty, J. D., McCallum, A., and Pereira, F. C. N. (2001). Conditional random fields: Probabilistic models for segmenting and labeling sequence data. In Proceedings of the Eighteenth International Conference on Machine Learning, ICML ’01, pages 282–289, San Francisco, CA, USA. Morgan Kaufmann Publishers Inc.

Lample, G., Ballesteros, M., Subramanian, S., Kawakami, K., and Dyer, C. (2016). Neural architectures for named entity recognition. In Proceedings of the 2016 Conference of the North American Chapter of the Association for Computational Linguistics: Human Language Technologies, pages 260–270. Association for Computational Linguistics.

Leaman, R., Miller, C., and Gonzalez, G. (2009). Enabling recognition of diseases in biomedical text with machine learning: corpus and benchmark. In Proceedings of the 2009 Symposium on Languages in Biology and Medicine, volume 82.

Lee, J. Y., Dernoncourt, F., and Szolovits, P. (2017). Transfer learning for named-entity recognition with neural networks. CoRR, abs/1705.06273.

Leser, U. and Hakenberg, J. (2005). What makes a gene name? named entity recognition in the biomedical literature. Briefings in bioinformatics, 6(4), 357–369.

Li, G., Ross, K. E., Arighi, C. N., Peng, Y., Wu, C. H., and Vijay-Shanker, K. (2015). miRTex: A text mining system for miRNA-gene relation extraction. PLOS Computational Biology, 11(9), e1004391.

Li, J., Sun, Y., Johnson, R. J., Sciaky, D., Wei, C.-H., Leaman, R., Davis, A. P., Mattingly, C. J., Wiegers, T. C., and Lu, Z. (2016). Biocreative v cdr task corpus: a resource for chemical disease relation extraction. Database, 2016, baw068.

Li, Q. (2012). Literature survey: domain adaptation algorithms for natural language processing. Department of Computer Science The Graduate Center, The City University of New York, pages 8–10.

Ling, W., Luís, T., Marujo, L., Astudillo, R. F., Amir, S., Dyer, C., Black, A. W., and Trancoso, I. (2015). Finding function in form: Compositional character models for open vocabulary word representation. arXiv preprint arXiv:1508.02096.

Mikolov, T., Sutskever, I., Chen, K., Corrado, G. S., and Dean, J. (2013). Distributed representations of words and phrases and their compositionality. In Advances in neural information processing systems, pages 3111–3119.

Mou, L., Meng, Z., Yan, R., Li, G., Xu, Y., Zhang, L., and Jin, Z. (2016). How transferable are neural networks in nlp applications? arXiv preprint arXiv:1603.06111.

Neves, M., Damaschun, A., Kurtz, A., and Leser, U. (2012). Annotating and evaluating text for stem cell research. In Proceedings of the Third Workshop on Building and Evaluation Resources for Biomedical Text Mining (BioTxtM 2012) at Language Resources and Evaluation (LREC). Istanbul, Turkey, pages 16–23.

Oquab, M., Bottou, L., Laptev, I., and Sivic, J. (2014). Learning and transferring mid-level image representations using convolutional neural networks. In Proceedings of the 2014 IEEE Conference on Computer Vision and Pattern Recognition, CVPR ’14, pages 1717–1724, Washington, DC, USA. IEEE Computer Society.

Pafilis, E., Frankild, S. P., Fanini, L., Faulwetter, S., Pavloudi, C., Vasileiadou, A., Arvanitidis, C., and Jensen, L. J. (2013). The SPECIES and ORGANISMS resources for fast and accurate identification of taxonomic names in text. PLoS ONE, 8(6), e65390.

Pan, S. J. and Yang, Q. (2010). A survey on transfer learning. IEEE Transactions on knowledge and data engineering, 22(10), 1345–1359.

Pennington, J., Socher, R., and Manning, C. (2014). Glove: Global vectors for word representation. In Proceedings of the 2014 conference on empirical methods in natural language processing (EMNLP), pages 1532–1543.

Pyysalo, S., Ginter, F., Heimonen, J., Bjorne, J., Boberg, J., Jarvinen, J., and Salakoski, T. (2007). Bioinfer: a corpus for information extraction in the biomedical domain. BMC Bioinformatics, 8(1), 50.

Pyysalo, S., Ginter, F., Moen, H., Salakoski, T., and Ananiadou, S. (2013). Distributional semantics resources for biomedical text processing. In Proceedings of the 5th International Symposium on Languages in Biology and Medicine, Tokyo, Japan. LBM.

Rastegar-Mojarad, M., Ye, Z., Kolesar, J. M., Hebbring, S. J., and Lin, S. M. (2015). Opportunities for drug repositioning from phenome-wide association studies. Nature biotechnology, 33(4), 342.

Rebholz-Schuhmann, D., Yepes, A. J. J., van Mulligen, E., Kang, N., Kors, J., Milward, D., Corbett, M., Buyko, E., Beisswanger, E., and Hahn, U. (2010). Calbc silver standard corpus. Journal of Bioinformatics and Computational Biology, 8(1), 163–179.

Reimers, N. and Gurevych, I. (2017). Optimal hyperparameters for deep lstm-networks for sequence labeling tasks. arXiv preprint arXiv:1707.06799.

Smith, L., Tanabe, L. K., Ando, R. J. n., Kuo, C.-J., Chung, I.-F., Hsu, C.-N., Lin, Y.-S., Klinger, R., Friedrich, C. M., Ganchev, K., Torii, M., Liu, H., Haddow, B., Struble, C. A., Povinelli, R. J., Vlachos, A., Baumgartner, W. A., Hunter, L., Carpenter, B., Tsai, R. T.-H., Dai, H.-J., Liu, F., Chen, Y., Sun, C., Katrenko, S., Adriaans, P., Blaschke, C., Torres, R., Neves, M., Nakov, P., Divoli, A., Maña-López, M., Mata, J., and Wilbur, W. J. (2008). Overview of biocreative ii gene mention recognition. Genome Biology, 9(2), S2.

Stamate, C., Magoulas, G. D., and Thomas, M. S. (2015). Transfer learning approach for financial applications. arXiv preprint arXiv:1509.02807.

Thrun, S. (1996). Is learning the n-th thing any easier than learning the first? In Advances in neural information processing systems, pages 640–646.

Verspoor, K., Jimeno Yepes, A., Cavedon, L., McIntosh, T., Herten-Crabb, A., Thomas, Z., and Plazzer, J.-P. (2013). Annotating the biomedical literature for the human variome. Database, 2013, bat019.

Wang, D. and Zheng, T. F. (2015). Transfer learning for speech and language processing. In Signal and Information Processing Association Annual Summit and Conference (APSIPA), 2015 Asia-Pacific, pages 1225–1237. IEEE.

Wang, X., Tsujii, J., and Ananiadou, S. (2010). Disambiguating the species of biomedical named entities using natural language parsers. Bioinformatics, 26(5), 661–667.

Wang, Z.-Y. and Zhang, H.-Y. (2013). Rational drug repositioning by medical genetics. Nature Biotechnology, 31(12), 1080–1082.

Weiss, K., Khoshgoftaar, T. M., and Wang, D. (2016). A survey of transfer learning. Journal of Big Data, 3(1).

Yosinski, J., Clune, J., Bengio, Y., and Lipson, H. (2014). How transferable are features in deep neural networks? CoRR, abs/1411.1792.

Young, S. R., Rose, D. C., Karnowski, T. P., Lim, S.-H., and Patton, R. M. (2015). Optimizing deep learning hyper-parameters through an evolutionary algorithm. In Proceedings of the Workshop on Machine Learning in High-Performance Computing Environments, MLHPC ’15, pages 4:1–4:5, New York, NY, USA. ACM.

Zeiler, M. D. and Fergus, R. (2013). Visualizing and understanding convolutional networks. CoRR, bs/1311.2901.

Zhou, X., Menche, J., Barabási, A.-L., and Sharma, A. (2014). Human symptoms–disease network. Nature Communications, 5.

